# Integrated aqueous humor ceRNA and miRNA-TF-mRNA network analysis reveals potential molecular mechanisms governing primary open-angle glaucoma pathogenesis

**DOI:** 10.1101/2020.07.17.208397

**Authors:** Xiaoqin Wang, Ming Chen, Liuzhi Zeng, Longqian Liu

## Abstract

Primary open-angle glaucoma (POAG) is the leading cause of blindness globally, which develops through complex and poorly understood biological mechanisms. Herein, we conducted an integrated bioinformatics analysis of extant aqueous humor (AH) gene expression datasets in order to identify key genes and regulatory mechanisms governing POAG progression. We downloaded AH gene expression datasets (GSE101727 and GSE105269) corresponding to healthy controls and POAG patients from the Gene Expression Omnibus. We then identified mRNAs, microRNAs (miRNAs), and long non-coding RNAs (lncRNAs) that were differentially expressed (DE) between control and POAG patients. DEmRNAs and DElncRNAs were then subjected to pathway enrichment analyses, after which a protein-protein interaction (PPI) network was generated. This network was then expanded to establish lncRNA-miRNA-mRNA and miRNA-transcription factor(TF)-mRNA networks. In total, the GSE101727 dataset was used to identify 2746 DElncRNAs and 2208 DEmRNAs, while the GSE105269 dataset was used to identify 45 DEmiRNAs. We ultimately constructed a competing endogenous RNA (ceRNA) network incorporating 37, 5, and 14 of these lncRNAs, miRNAs and mRNAs, respectively. The proteins encoded by these 14 hub mRNAs were found to be significantly enriched for activities that may be linked to POAG pathogenesis. In addition, we generated a miRNA-TF-mRNA regulatory network containing 2 miRNAs (miR-135a-5p and miR-139-5p), 5 TFs (TGIF2, TBX5, HNF1A, TCF3, and FOS) and 5 mRNAs (SHISA7, ST6GAC2, TXNIP, FOS, and DCBLD2). The SHISA7, ST6GAC2, TXNIP, FOS, and DCBLD2 genes that may be viable therapeutic targets for the prevention or treatment of POAG, and regulated by the TFs (TGIF2, HNF1A, TCF3, and FOS).

## Introduction

Primary open-angle glaucoma (POAG) can result in increased intraocular pressure (IOP), irreversible damage to the optic nerve, and eventually blindness, and it is the leading cause of such blindness globally[1,2]. As of 2013, an estimated 64.3 million people between the ages of 40 and 80 were estimated to be affected by glaucoma worldwide, with this number being forecast to rise to 76.0 million and 111.8 million in 2020 and 2040, respectively[3]. Elevated IOP is known to be a major risk factor for POAG development that is normally regulated by homeostatic aqueous humor (AH) production and outflow[4]. Hundreds of different genes are believed to govern POAG development and progression. Indeed, as this disease develops through complex and poorly understood biological mechanisms, few effective biomarkers of POAG have been identified to date. While several studies have explored the roles of specific genes or transcriptional regulators (TFs) in the context of POAG incidence, the full complexity of the interactions between long non-coding RNAs (lncRNAs), mRNAs, microRNAs (miRNAs) and TFs in this disease context remains to be fully elucidated.

The recently proposed competing endogenous RNA (ceRNA) hypothesis suggests that specific RNAs can interact with one another through complementary miRNA response elements (MREs), enabling certain RNAs to function as molecular ‘sponges’ capable of sequestering and thereby modulating the functionality of specific target miRNAs[5]. Such a model enables researchers to develop putative ceRNA networks associating the expression of specific mRNAs with the activity of other non-coding RNA types including miRNAs, lncRNAs, and circular RNAs (circRNAs)[6]. TFs and miRNAs serve as essential regulators of mRNA expression at the transcriptional and post-transcriptional stages, respectively, making them vital in both physiological and pathological contexts. These regulatory processes, however, do not happen in isolation, and as such TFs and miRNAs have the potential to impact one another and to influence target mRNA expression in a complex and dynamic fashion that has yet to be studied in detail in the context of POAG development and progression.

Herein, we performed an integrative analysis of two POAG-related Gene Expression Omnibus (GEO) datasets in order to identify lncRNAs, miRNAs and mRNAs that were differentially expressed (DE) in the AH of POAG patients relative to healthy controls. We then utilized the gene ontology (GO) and Kyoto encyclopedia of genes and genomes (KEGG) databases to conduct functional enrichment analyses of identified DE genes (DEGs), after which a protein-protein interaction (PPI) network was constructed. We further conducted systematic integrated analyses of identified DElncRNAs, DEmiRNAs, and DEmRNAs based on their co-expression profiles in light of the ceRNA hypothesis, enabling us to conduct successive lncRNA-miRNA-mRNA and miRNA-TF-mRNA network analyses. The overall goal of these bioinformatics analyses was to identify genes related to POAG incidence and to provide novel insight into the molecular mechanisms governing this debilitating disease.

## Materials and methods

### Selection of datasets

Two human AH microarray datasets were downloaded from GEO (http://www.ncbi.nlm.nih.gov/geo/) for use in this study, including GSE101727, which included lncRNA and mRNA expression profiles, and GSE105269, which included miRNA expression profiles. The GSE101727 dataset was based upon the GPL21827 platform (Agilent-079487 Arraystar Human LncRNA microarray V4) and contained 10 control and 10 POAG samples[7], while the GSE105269 dataset was based upon the GPL24158 platform (NanoString nCounter Human v3 miRNA Assay (NS_H_miR_v3a)) and included 11 control and 12 POAG samples[8].

### Identification of DEGs

Prior to DEG identification, dataset quality was evaluated through the use of box plots and principal component analyses. The linear models for microarray data (LIMMA) and Affy R packages were then used for differential expression analyses, with *P* < 0.05 and |log2 fold change (FC)| ≥ 1 as the cut-off criteria for DEG identification. This strategy was used to identify DElncRNAs, DEmiRNAs, and DEmRNAs, which were then arranged in heat maps and subjected to downstream analyses.

### Functional enrichment analyses

The GO (http://www.geneontology.org) functions of identified DEGs, including enriched biological processes (BPs), molecular functions (MFs), and cellular components (CCs), were identified based upon available annotation to explore the functional roles of these proteins. KEGG (http://www.genome.jp/kegg/) analyses were similarly used to evaluate the pathways in which these DEGs are functionally enriched in the context of POAG. The R Clusterprofiler package was used to conduct GO and KEGG analyses, with P ≤ 0.05 as the threshold of significance.

### ceRNA network construction

A ceRNA network was constructed by first evaluating the putative interactions between DElncRNAs, DEmiRNAs and DEmRNAs that were differentially regulated in POAG patient samples. Interactions between DElncRNAs and DEmiRNAs were predicted using miRcode, after which miRanda, miRMap, miTarBase, miRDB, and Targetscan were employed to identify predicted relationships between DEmiRNAs and DEmRNAs of interest. These predicted interactions were then used to construct a final ceRNA network.

### PPI network construction

The Search Tool for the Retrieval of Interacting Genes (STRING, https://string-db.org/cgi/input.pl) database was utilized for PPI network construction, with the resultant network being visualized using Cytoscape 3.4.0.

### miRNA-TF-mRNA regulatory network construction

TF binding sites were initially predicted with the TFBS Tools R package, enabling us to predict TFs likely to regulate the transcription of 14 key POAG-related mRNAs present within our ceRNA network. Experimentally validated databases (miRDB, TargetScan, miRanda, miRMap and miTarBase) were then employed to predict miRNA-TF and miRNA-mRNA interactions, allowing us to construct a miRNA-TF-mRNA regulatory network that was visualized using Cytoscape 3.4.0.

## Results

### Differentially expressed lncRNA, miRNA, and mRNA identification

In the GSE101727 dataset, we identified 2746 DElncRNAs (1399 upregulated and 1347 downregulated) and 2208 DE mRNAs (1469 upregulated and 739 down regulated) when comparing POAG patient samples to control samples (P < 0.05 and |log2 FC| ≥ 2.0) (Table 1). These DElncRNAs and DEmRNAs were arranged in heat maps and volcano plots for ease of visualization (Fig. 1A and 1B). Similar analyses of the GSE105269 dataset led us to identify 45 DEmiRNAs (23 upregulated and 22 downregulated) in POAG samples relative to normal control samples (P < 0.05 and |log2 FC| ≥ 1.2) (Table 2). These DEmiRNAs were additionally organized into volcano plots and heat maps (Fig. 1C and 1D). For further details regarding these DEG analyses, see File S1.

**Fig. 1.**
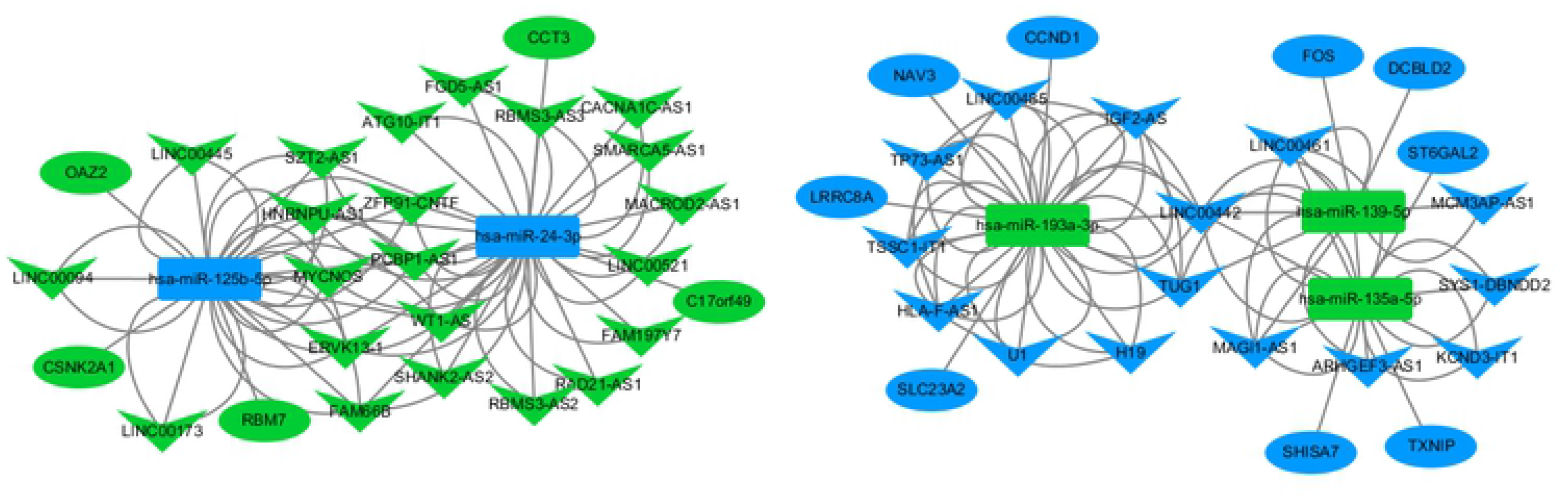
Differentially expressed lncRNAs, mRNAs, and miRNAs identified by comparing control and POAG patients in the GSE101727 (A) and GSE105269 (B) datasets.

**Table 1.**
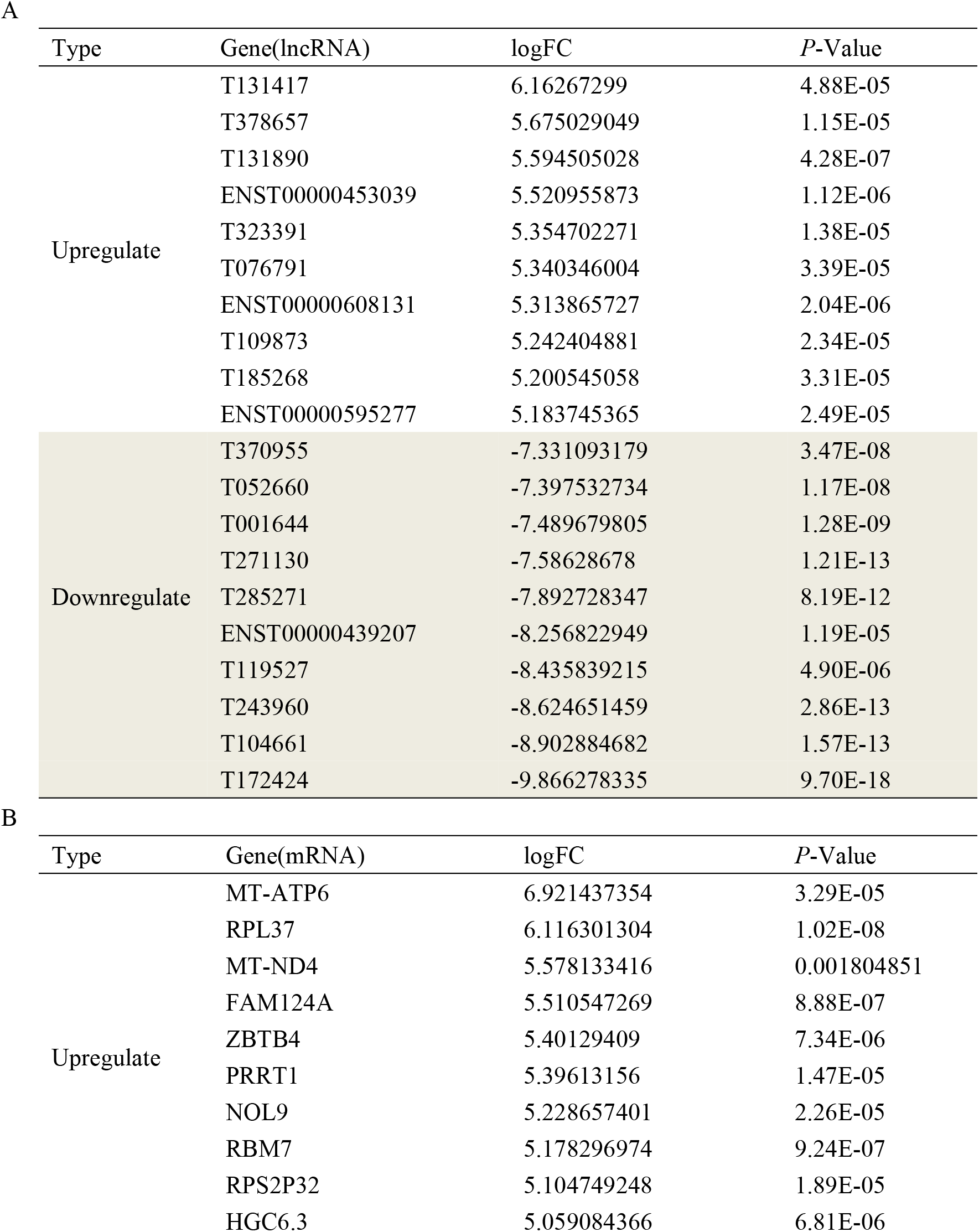
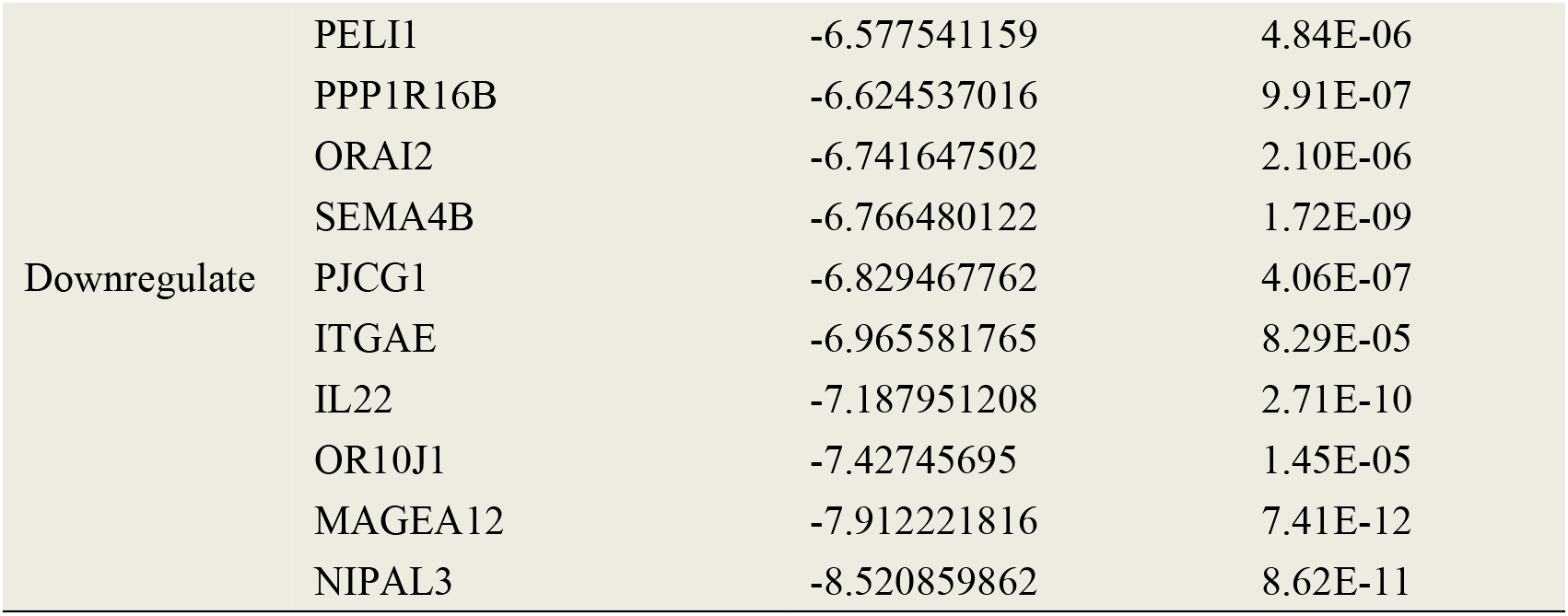
The top 20 differentially expressed lncRNAs (A) and mRNAs (B) in the AH when comparing POAG patient and control samples in the GSE101727 dataset

**Table 2.** The top 20 differentially expressed miRNAs in the AH when comparing POAG patient samples and control samples in the GSE105269 dataset

| Type | Gene(miRNA) | logFC | P-Value |
| --- | --- | --- | --- |
| Upregulate | hsa-miR-451a | 1.099764108 | 0.032177995 |
|  | hsa-miR-3140-5p | 0.7593001 | 0.034883538 |
|  | hsa-miR-3190-3p | 0.660045773 | 0.017180336 |
|  | hsa-miR-194-5p | 0.657009272 | 0.020429938 |
|  | hsa-miR-135a-5p | 0.642775355 | 0.045576341 |
|  | hsa-miR-320e | 0.596668708 | 0.024266619 |
|  | hsa-miR-648 | 0.5066355 | 0.007511757 |
|  | hsa-miR-137 | 0.497752609 | 0.049674124 |
|  | hsa-miR-224-5p | 0.497159513 | 0.04279984 |
|  | hsa-miR-1244 | 0.482940146 | 0.007688872 |
| Downregulate | hsa-miR-484 | -0.505411455 | 0.034413007 |
|  | hsa-miR-431-5p | -0.50830537 | 0.018242336 |
|  | hsa-miR-1910-5p | -0.534734696 | 0.032714754 |
|  | hsa-miR-2682-5p | -0.537553037 | 0.006452898 |
|  | hsa-miR-517c-3p | -0.54289098 | 0.000659242 |
|  | hsa-miR-375 | -0.545806863 | 0.004278049 |
|  | hsa-miR-106b-5p | -0.579127165 | 0.035851945 |
|  | hsa-miR-3195 | -0.599964555 | 0.031150973 |
|  | hsa-miR-125b-5p | -0.630025636 | 0.009991458 |
|  | hsa-miR-204-5p | -0.844884632 | 0.018554867 |

### Functional enrichment analyses

We next conducted GO and KEGG analyses to explore the potential functional roles of these identified DElncRNAs and DEmRNAs in the context of POAG. These DEGs were enriched in GO terms associated with nuclear-transcribed mRNA catabolic processes, nonsense-mediated decay, translational initiation, cytosolic ribosomes, ribonucleoprotein complexes, and structural constituents of ribosomes (Fig. 2A-C; Table 3). KEGG pathway analyses similarly revealed these DEGs to be enriched in pathways associated with ribosomes, spliceosomes, and the proteasome (Fig. 2D; Table 4). For full details regarding these functional enrichment analyses, see File S2.

**Fig. 2.**
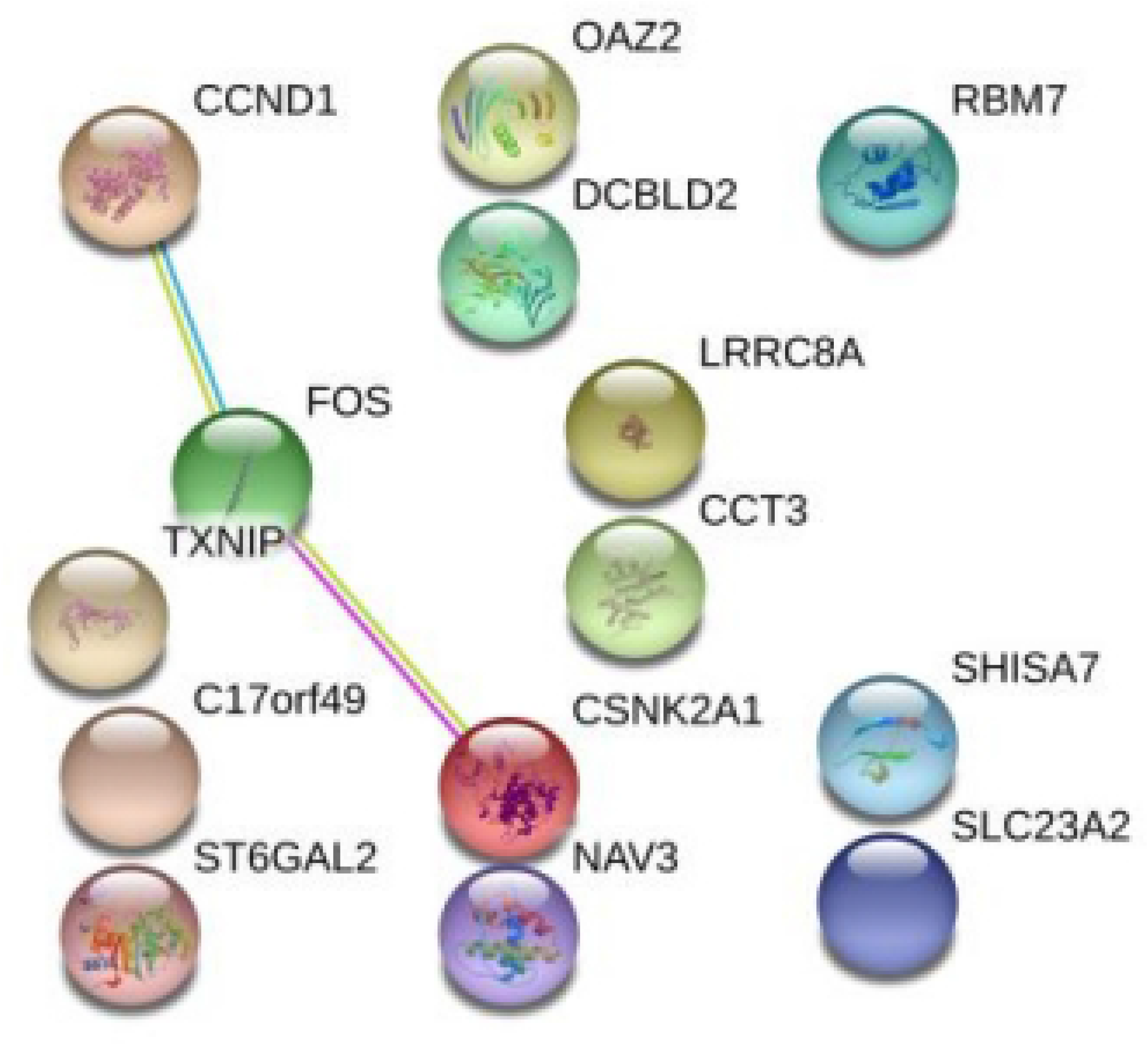
The top 10 GO and KEGG terms associated with DElncRNAs and DEmRNAs. (A) Biological processes (BPs); (B) Cellular components (CCs); (C) Molecular functions (MFs). (D) Enriched KEGG pathways. In all figures, circle size is proportional to gene number, while circle color corresponds to the adjusted P-value, with red corresponding to a smaller adjusted P-value.

**Table 3.**
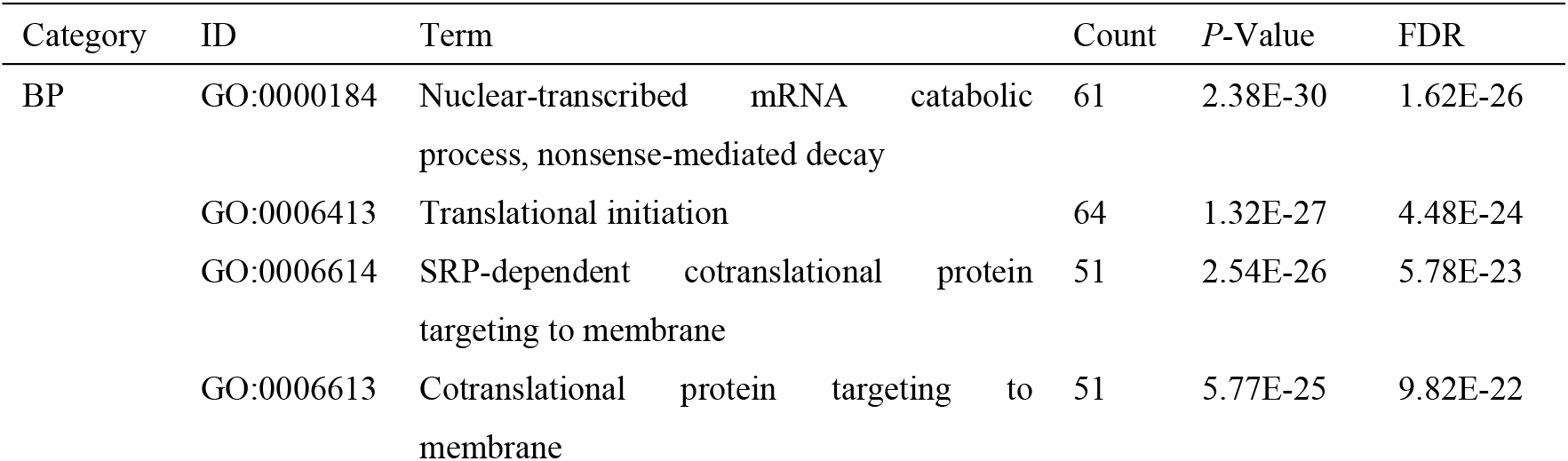
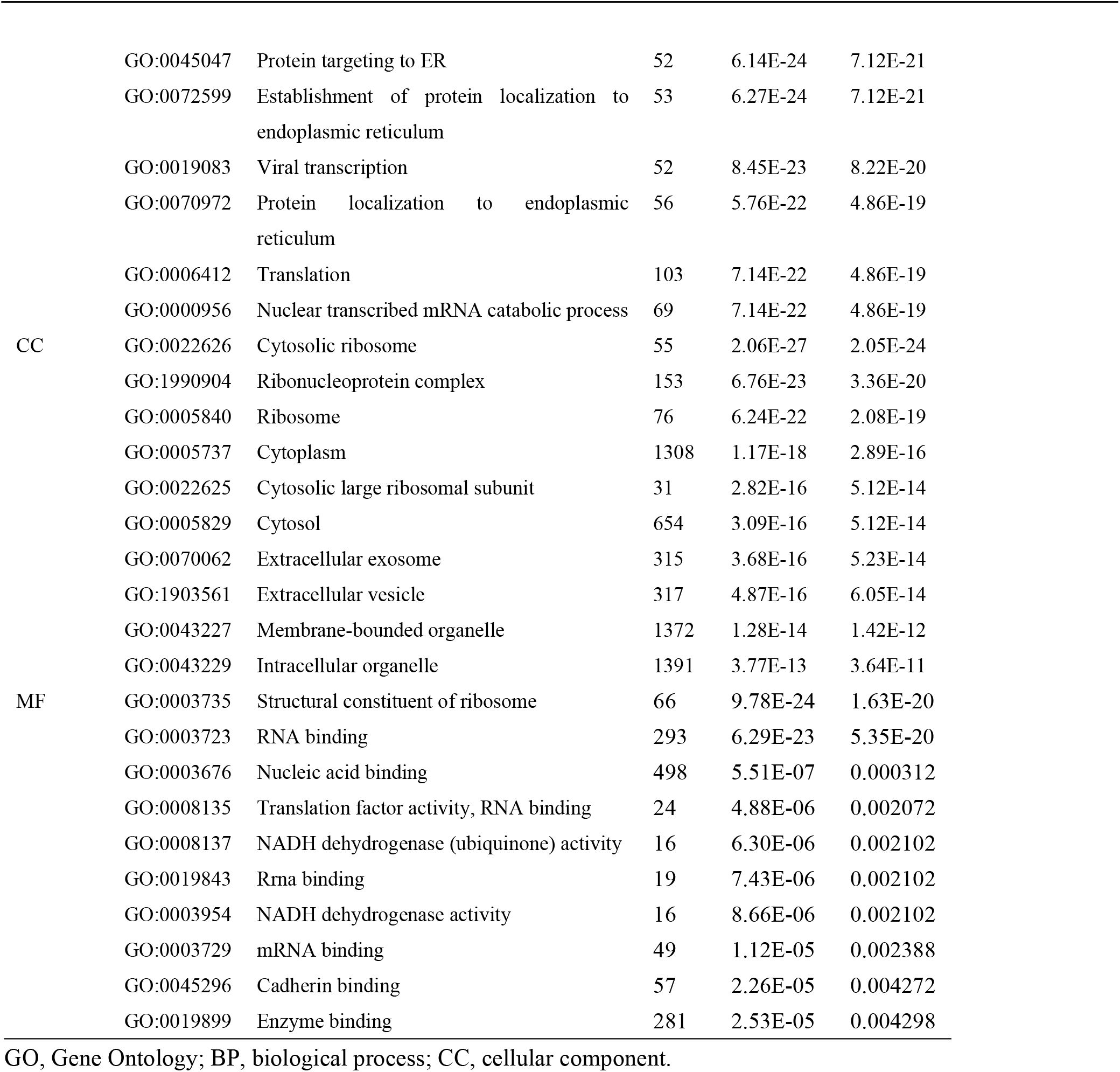
The top 10 significantly enriched GO terms associated with differentially expressed lncRNAs and mRNAs

**Table 4.**
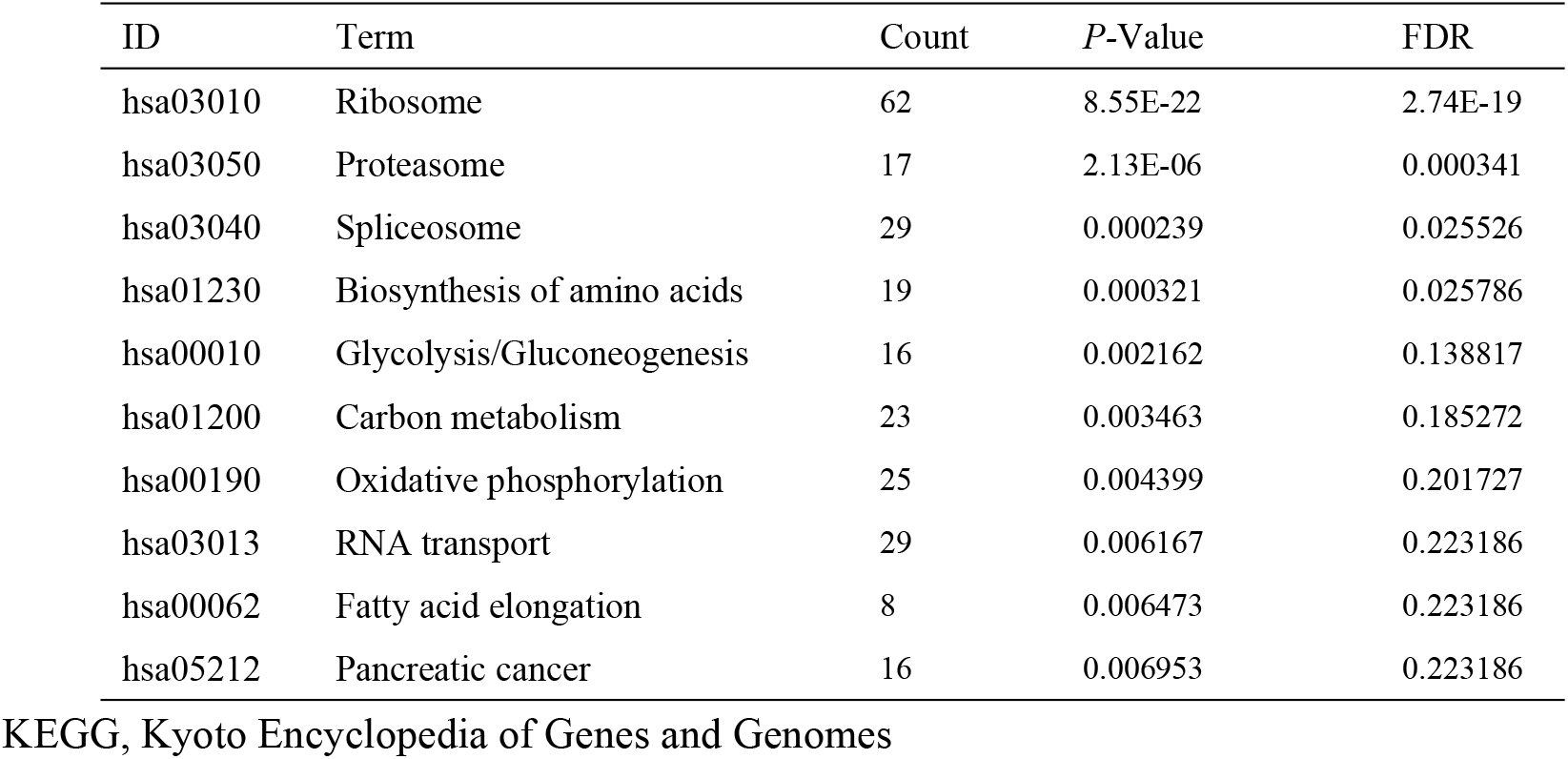
The top 10 significantly enriched KEGG pathways associated with differentially expressed lncRNAs and mRNAs

| ID | Term | Count | P-Value | FDR |
| --- | --- | --- | --- | --- |
| hsa03010 | Ribosome | 62 | 8.55E-22 | 2.74E-19 |
| hsa03050 | Proteasome | 17 | 2.13E-06 | 0.000341 |
| hsa03040 | Spliceosome | 29 | 0.000239 | 0.025526 |
| hsa01230 | Biosynthesis of amino acids | 19 | 0.000321 | 0.025786 |
| hsa00010 | Glycolysis/Gluconeogenesis | 16 | 0.002162 | 0.138817 |
| hsa01200 | Carbon metabolism | 23 | 0.003463 | 0.185272 |
| hsa00190 | Oxidative phosphorylation | 25 | 0.004399 | 0.201727 |
| hsa03013 | RNA transport | 29 | 0.006167 | 0.223186 |
| hsa00062 | Fatty acid elongation | 8 | 0.006473 | 0.223186 |
| hsa05212 | Pancreatic cancer | 16 | 0.006953 | 0.223186 |

KEGG, Kyoto Encyclopedia of Genes and Genomes

### Construction of a predictive ceRNA network

A putative lncRNA-miRNA-mRNA ceRNA network was next constructed by identifying lncRNAs and mRNAs targeted by miRNAs in the GSE101727 dataset (File S3). A total of 55 DElncRNAs in this study were predicted to interact with 8 miRNAs, while 604 mRNAs were predicted to be targets of 6 identified DEmiRNAs. By integrating these predictive analyses, we were able to generate a network of putative interactions among 37 DElncRNAs, 5 DEmiRNAs, and 14 DEmRNAs (Fig. 3; Table 5).

**Fig. 3.**
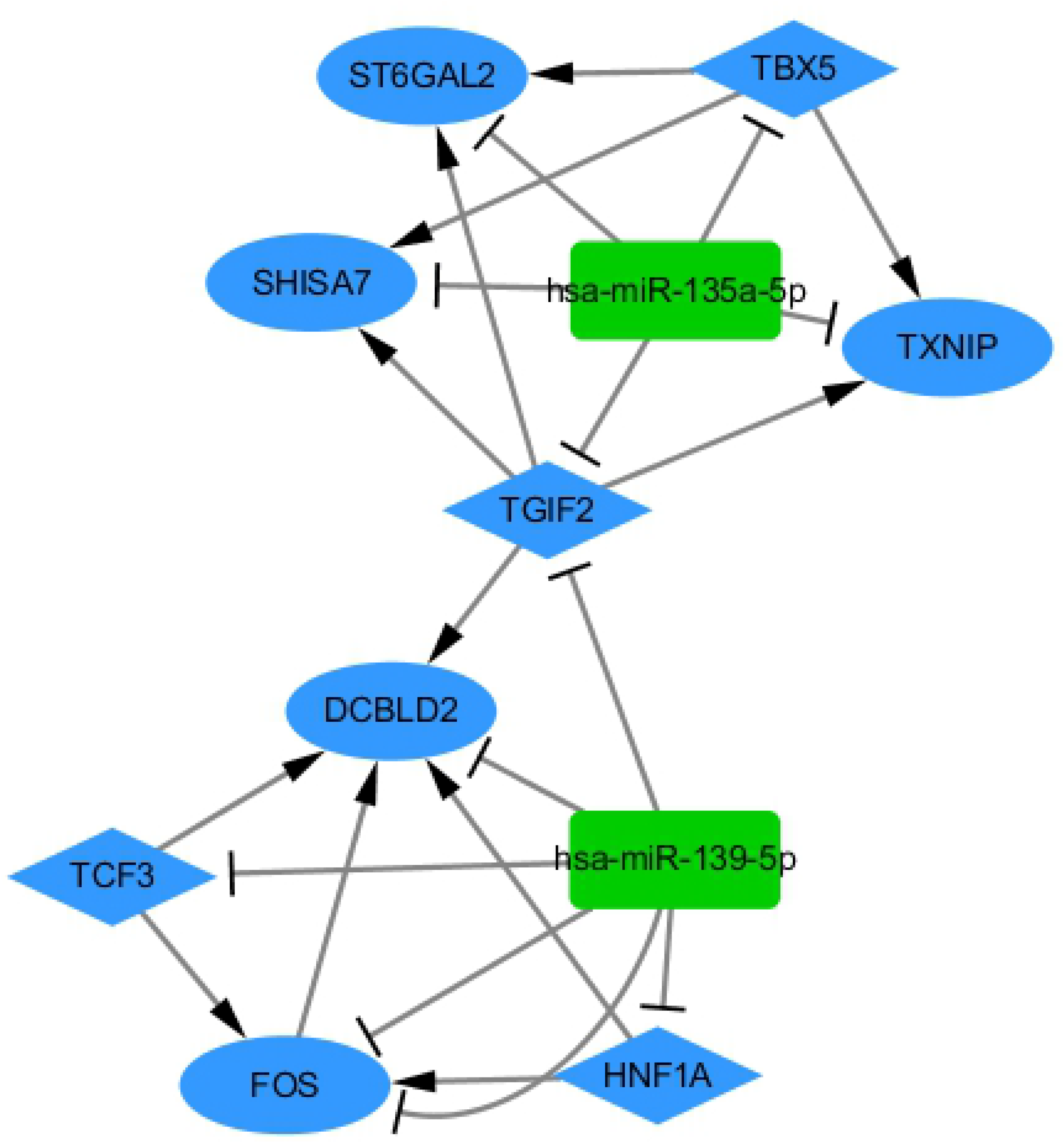
The POAG lncRNA-miRNA-mRNA ceRNA interaction network. Triangles, rectangles, and ovals correspond to lncRNAs, miRNAs, and mRNAs, respectively, with blue and green corresponding to downregulation and upregulation, respectively. Gray edges indicate interactions between RNAs. ceRNA, competing endogenous RNA; DElncRNAs, differentially expressed lncRNAs; DEmiRNAs, differentially expressed miRNAs; DEmRNAs, differentially expressed mRNAs.

**Table 5.**
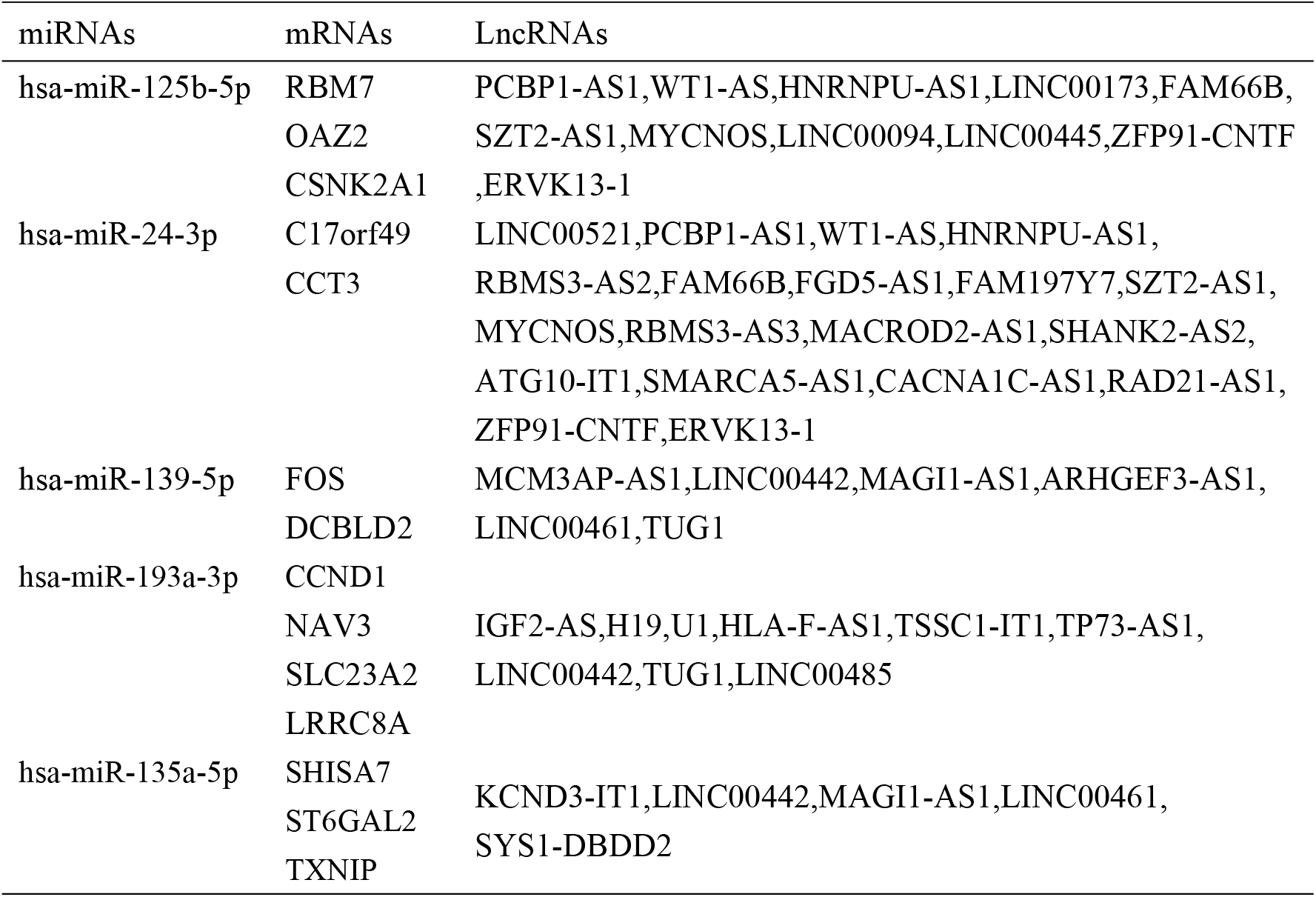
Representative interactions among lncRNAs, miRNAs, and mRNAs

### PPI network construction

A PPI network incorporating the 14 mRNAs identified in our ceRNA network was next constructed (Fig. 4). This analysis revealed that a subset of these proteins were predicted to interact with one another, suggesting that these key proteins may play interrelated roles in the context of POAG pathogenesis.

**Fig. 4.**
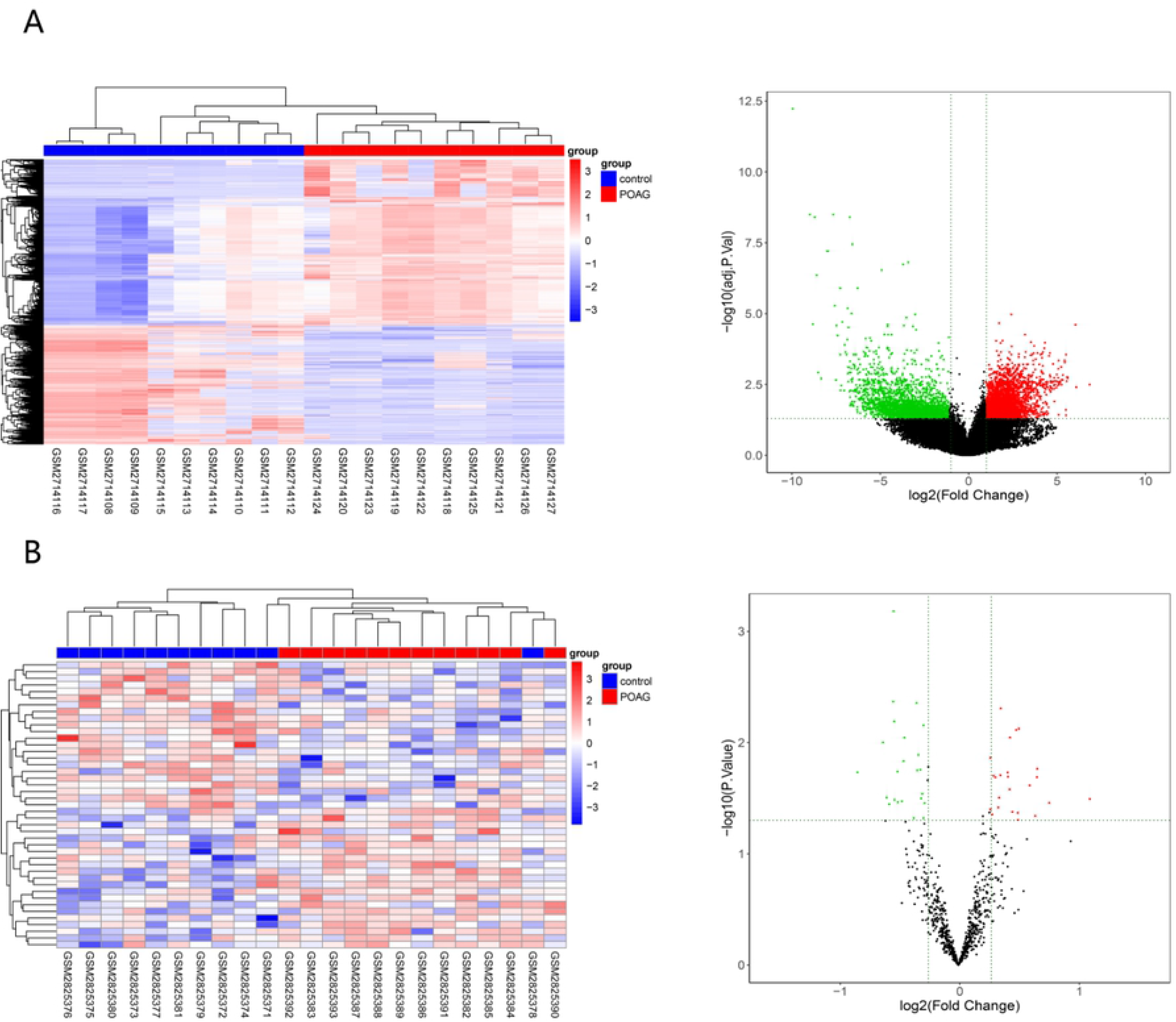
A PPI network of POAG-associated ceRNA network-related DEmRNAs. The 3D structure of the indicated proteins is represented within nodes, when known. Edges correspond to protein-protein interactions, with blue, purple, and yellow edges corresponding to interactions identified based upon curated databases, experimental validation, and text mining, respectively.

### miRNA-TF-mRNA regulatory network construction

In our final analyses, we identified 383 putative TF binding associations and 6 TFs predicted to regulate the expression of these 14 POAG-related hub RNAs. Integrated miRNA, TF, and mRNA pairs are compiled in Table 6, and a predicted miRNA-TF-mRNA regulatory network was developed based upon these analyses (Fig. 5). This network incorporated 2 miRNAs (miR-135a-5p and miR-139-5p), 5 TFs (TGIF2, TBX5, HNF1A, TCF3, and FOS) and 5 mRNAs (SHISA7, ST6GAC2, TXNIP, FOS, and DCBLD2). We additionally determined that miR-135a-5p was predicted to inhibit 2 TFs (TGIF2 and TBX5) and 3 mRNAs (SHISA7, ST6GAC2, and TXNIP), while miR-139-5p was predicted to inhibit 4 TFs (TGIF2, HNF1A, TCF3, and FOS) and 2 mRNAs (FOS and DCBLD2). These findings suggested that different miRNAs were able to co-regulate the expression of specific genes, indicating that these miRNAs may play complex and interrelated roles in the context of POAG incidence. Of the 5 TFs identified in these analyses, TGIF2 was predicted to regulate the expression of four mRNAs (ST6GAL2, SHISA7, TXNIP, and DCBLD2). In addition, FOS was predicted to serve as a regulator of DCBLD2, which was also co-regulated by TCF3 and HNF1A in our regulatory network. Similarly, DCBLD2 was co-regulated by TGIF2, HNF1A, TCF3, and FOS.

**Table 6.**
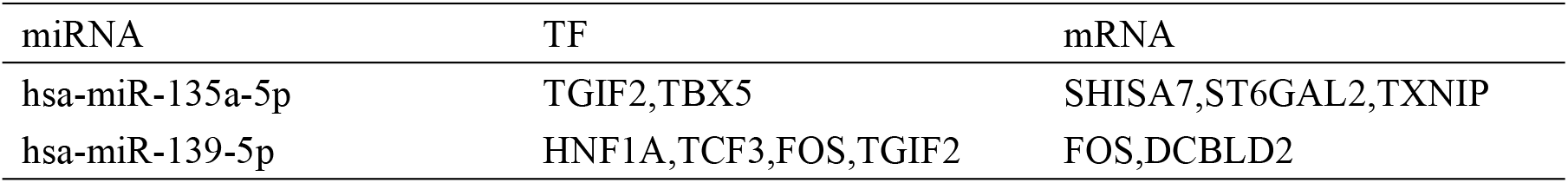
Integrated miRNA, TF and mRNA pairs

**Fig. 5.**
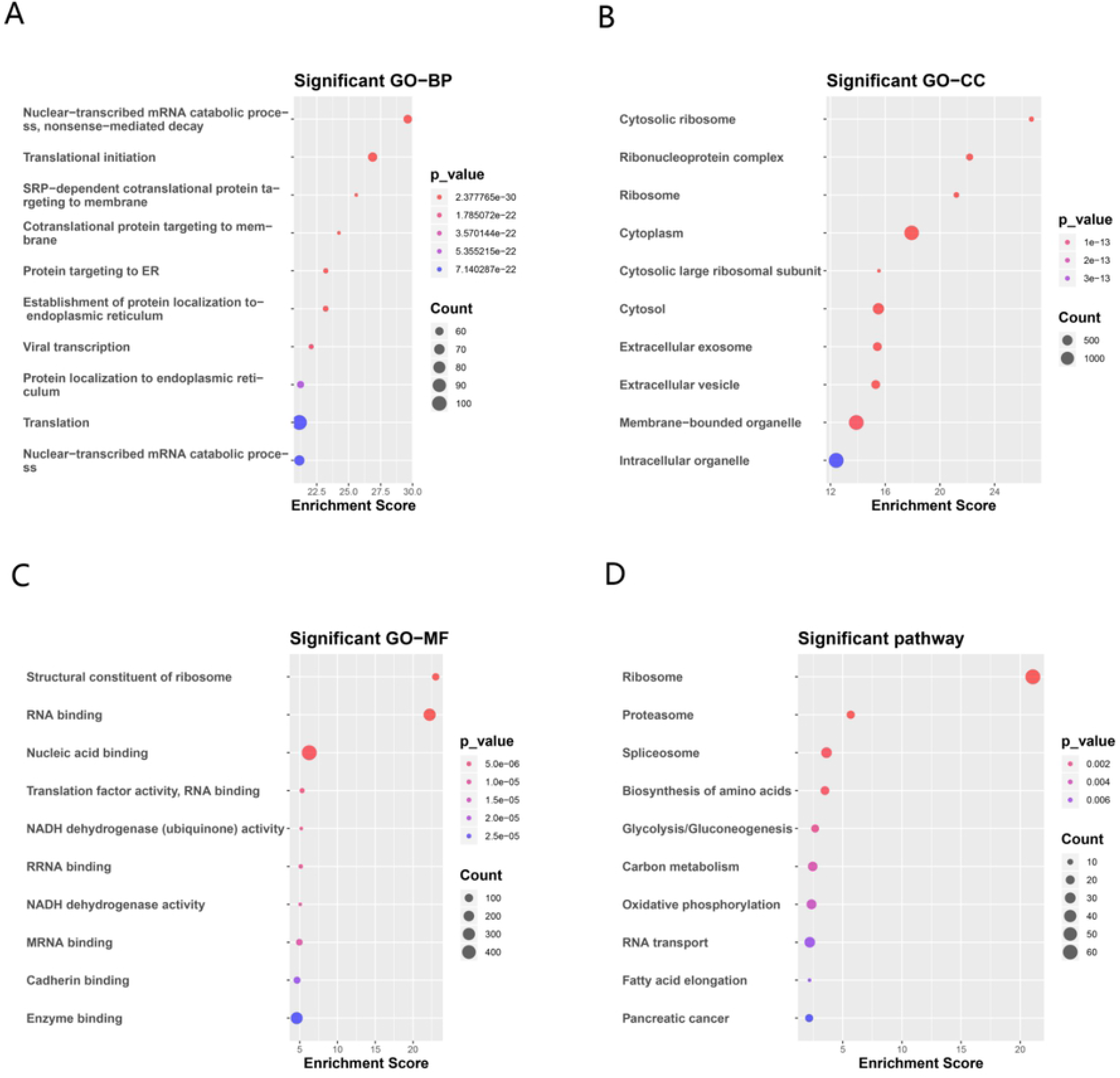
miRNA-TF-mRNA regulatory network. Diamonds, rectangles, and ovals correspond to TFs, miRNAs, and mRNAs, respectively, with blue and green indicating downregulation and upregulation, respectively. Arrows indicate activation, with other lines corresponding to repression. miRNA, micro RNA; TF, transcription factor; mRNA, message RNA.

## Discussion

Elevated IOP can result in axonal compression of the optic nerve and the apoptotic death of retinal ganglion cells (RGCs), thereby impairing the normal homeostatic balance between AH production and outflow within the anterior chamber[9,10]. Given the pathological nature of such dysregulation, molecular mechanisms regulating the AH and IOP can influence glaucoma development, and prior studies have identified multiple pathways and individual genes associated with POAG incidence[11,12]. Few analyses to date, however, have conducted systematic analyses to identify relationships between lncRNAs, miRNAs, TF, and mRNAs in this pathological context. Herein, we therefore conducted a comprehensive integrative bioinformatics analysis of the molecular basis for POAG pathogenesis, leading us to identify multiple genes and TFs that are potentially associated with this condition. By analyzing two extant microarray datasets, we were able to identify 2746 DElncRNAs and 2208 DEmRNAs in the GSE101727 dataset and 45 DEmiRNAs in the GSE105269 dataset. Using GO and KEGG pathway analyses, we further explored the potential pathogenic roles of these lncRNAs and mRNAs, and we thereafter constructed lncRNA-miRNA-mRNA and miRNA-TF-mRNA networks to evaluate their interrelated regulatory functions.

In eukaryotes, both TFs and miRNAs serve as primary regulators of gene expression, with TFs serving to directly control DNA transcription[13,14] and miRNAs functioning by binding to conserved MREs to disrupt target mRNA translation at the post-transcriptional level[15,16,17,18]. In prior analyses, miRNAs have been shown to control anterior chamber shape, IOP, and RGC apoptosis via the regulation of specific target genes[19]. In addition, there is evidence that miRNAs can regulate diverse biochemical pathways in the context of glaucoma[20], and miRNAs including miR-125b-5p have been detected at elevated levels in human AH relative to human serum [21,22]. At a functional level, miR-139 has been found to regulate the WNT signaling pathway, which is a key mediator of TGFβ1-induced fibrosis[23]. As TGFβ signaling served to regulate the AH environment by regulating ocular hypertensive mediator gene expression[24], this suggests that this pathway may be associated with POAG incidence. We found that miR-125b-5p and mir-24-3p were downregulated in our ceRNA network, whereas miR-135a-5p, mir-193a-3p, and miR-139-5p were upregulated. Both miR-135a-5p and miR-139-3p can suppress the expression of TGIF2, which is a transcriptional regulator that is important in the context of cellular differentiation, proliferation, and embryonic development[25,26]. Herein, we identified TGIF2 as a direct miR-135a-5p and miR-139-5p target in the context of POAG, suggesting that the ability of these two miRNAs to modulate POAG progression may be a result of their ability to control TGIF2 expression.

FOS (c-Fos) is an inducible transcription factor and member of the activator protein 1(AP1) family[27].Through protein-protein interactions, FOS can additionally govern the transcriptional activity of other TFs[28,29]. We found that FOS was incorporated into our ceRNA and PPI networks, suggesting that it may be a key regulator of POAG development and progression. FOS expression is closely associated with the apoptotic death of various neuronal cell types[30,31], and there is also evidence that it plays a role in mediating RGC damage[32], underscoring its potential relevance to the etiology of POAG. TXNIP, which was first detected in 1, 25-dihydroxyvitamin D-3-treated HL-60 cells[33], can increase cellular sensitivity to oxidative stress and can drive bioenergetic imbalance, autophagy, and apoptotic cell death[34,35]. Oxidative stress is one of the primary drivers of IOP-associated RGC death, and TXNIP upregulation is observed in the context of RGC death induced by both optic nerve transection and elevated IOP[36,37]. When the expression of TXNIP is decreased, this is sufficient to inhibit RGC death[38]. RGC and optic nerve axonal degradation are the primary diverse of visual impairment and blindness in POAG patients. These data thus confirm that TXNIP is a key target in the context of POAG progression. Herein, we determined that miR-135a-5p and two TFs (TGIF2 and TBX5) were able to regulate TXNIP expression in the AH of POAG patients. DCBLD2 is a key regulator of cellular proliferation and vascular remodeling[39,40]. Our miRNA-TF-mRNA network suggested that HNF1A, TCF3, FOS and TGIF2 were able to stimulate DCBLD2 expression, indicating that DCBLD2 may influence POAG pathogenesis through these transcriptional mechanisms.

In summary, in the present study we identified a novel subset of genes that are differentially expressed in the AH of POAG patients relative to the AH of healthy controls, suggesting that these genes may be closely linked to the development and progression of this debilitating condition. Through our comprehensive bioinformatics analyses, we highlighted novel transcription and post-transcriptional regulatory mechanisms governing POAG incidence. We were further able to identify 5 genes (SHISA7, ST6GAC2, TXNIP, FOS, and DCBLD2) that may represent viable targets for the treatment or prevention of POAG, and that are regulated by the TFs TGIF2, HNF1A, TCF3, and FOS. While our findings provide new insight into the molecular etiology of POAG, further experimental validation of our results will be essential in order to confirm their relevance and to explore their therapeutic potential.

## ABBREVIATIONS

POAG: primary open-angle glaucoma
lncRNA: long non-coding RNA
mRNA: message RNA
miRNA: micro RNA
TF: transcription factor
IOP: intraocular pressure
AH: aqueous humor
GEO: Gene Expression Omnibus
GO: gene ontology
KEGG: Kyoto encyclopedia of genes and genomes
PPI: protein-protein interaction
BP: biological process
MF: molecular function
CC: cellular component

## Acknowledgements

We appreciate the assistance from Wenjing Xu with technical analysis.

## Supporting Information

Additional Supporting Information may be found in the online version of this article:

S1 File. Complete results of the DEGs analysis
S2 File. Complete results of the GO and KEGG analysis.
S3 File. miRNA targets

## Author Contributions

X.W., L.Z. and L.L. conceived and designed the experiments; X.W. conducted the bioinformatic analysis; X.W. wrote the original draft; L.Z., M.C. and L.L. revised the paper.

